# Cardio- and neurometabolic effects of lower-body pressure-supported exercise in obese non-diabetic women: Resetting autonomic imbalance?

**DOI:** 10.1101/202986

**Authors:** Ellen M. Godwin, Anthony D. Uglialoro, Andaleeb Ali, Leah Yearwood, Mary Ann Banerji, John G. Kral

**Affiliations:** Department of Physical Therapy, Long Island University, Brooklyn, New York, United States of America; Department of Orthopedics/Rehabilitation, SUNY Downstate Medical Center, Brooklyn, New York, United States of America; Department of Surgery, SUNY Downstate Medical Center, Brooklyn, New York, United States of America; Department of Medicine, SUNY Downstate Medical Center, Brooklyn, New York, United States of America

## Abstract

**Background:** Overnutrition and underactivity cause most chronic disease via inflammation and stress. Life-style changes such as diet is largely unsuccessful and exercise is painful, uncomfortable and difficult for people with diabesity, cardiorespiratory and joint diseases and cognitive decline affecting their ability to ambulate and adhere to exercise guidelines. Diets or exercise causing weight loss are stressful and trigger numerous redundant counter regulatory mechanisms defending lean body mass, explaining failures to sustain these behaviors. In this hypothesis-generating pilot study we used a NASA-developed weight supporting lower-body positive pressure (LBPP) treadmill providing comfortable low-amount, low intensity walking, challenging current exercise guidelines.

**Methods:** Sixteen nondiabetic, untrained, Black volunteer women (BMI 28-50), age 18-56 years were studied by anthropometry, analyses of energy expenditure and blood chemistry: oral glucose tolerance tests (OGTT) with insulin, C-peptide, GLP-1 and FFA and fasting lipids, cytokines, adipokines and appetitive peptides, before and after 10 weeks of twice weekly 30-minute weight supported LBPP treadmill sessions.

**Results:** We found novel baseline associations between gluco- and neuro-regulatory peptides and plasma lipids, inflammatory cytokines and appetitive hormones related to neurogenesis, mood and energy balance. Post-study, independent of body weight or energy expended there were significant decreases in OGTT plasma insulin (p=0.002) and GLP-1 (p=0.060) and fasting triglycerides (p=0.029), ghrelin (p=0.008) and changes in most molecules including increased leptin and beta-endorphin. Correlations between changes among different classes of peptides were highly significant, notably leptin - adiponectin, and beta-endorphin - oxytocin and orexin A. We propose synergy between low-amount, low-intensity exercise at levels below thresholds of increased sympathetic tone, and baro-physiological effects of LBPP normalizing parasympathetic tone.

**Conclusion:** Brief, low-dose, lower-body positive-pressure weight-supported treadmill exercise improved cardiometabolic fitness and exhibited favorable changes in neuro-regulatory peptides without weight loss in inner-city obese Black women.

## Introduction

Chronic human diseases, from atherosclerosis and neoplasia to organ failure are anthropogenic emanating from overnutrition and underactivity causing inflammation and stress and autonomic imbalance related to allostatic overload [1]. Increasing physical activity is easier than dieting and less stressful under normal circumstances. Exercise mitigates stress [2] and has beneficial effects on inflammatory pathways and numerous diseases [3], especially the inflammatory insulin resistant chronic overnutrition syndrome diabesity and its numerous diverse comorbidities. Both exercise and diabesity are pleiotropic, reciprocally affecting all body systems through similar pathways sharing gluco-regulatory, adipogenic, inflammatory, neurotrophic and appetitive molecules modulated by the parasympathetic nervous system [Supporting Table S1A].

Life-style change, the primary therapeutic and preventive intervention for numerous degenerative diseases includes diet and exercise. Extant recommendations and guidelines consonantly recommend 150 minutes of moderate-intensity activity, 75 minutes of vigorous-intensity activity, or some combination of moderate and vigorous activity with 2 days of resistance exercise weekly [27], but are rarely successful long enough to be effective. With the exception of early-life measures most interventions are implemented late in cumulative disease processes when the magnitude and rapidity of recommended interventions widely exceed those of cumulative causal factors, thus eliciting counter-regulatory non-homeostatic responses. Whereas most guidelines propose moderate to high intensity exercise to lose weight, there is little research investigating metabolic effects of lower levels of energy expenditure without weight loss [28, 29].

Our study tests the hypothesis that incremental increases in normative levels of natural forms of exertion in the “sedentary range” [30, 31], i.e. below an upper threshold that triggers compensatory sympatho-adrenal mechanisms, might reduce cardiometabolic risk sufficient to ameliorate or prevent diabesity in the absence of weight loss. We use a lower-body positive-pressure (LBPP) treadmill developed by the National Aeronautics and Space Administration (NASA) with a pressurized air chamber which offloads weight for easier and painless exercise [32] for rehabilitation of patients with orthopedic and neurological conditions [33, 34]. Our application is to enable ambulation in people with difficulties performing aerobic exercise using their largest muscle mass, viz. those with diabesity [35], a condition associated with sympathovagal imbalance [36]. The alternative, standing in water, supports body weight, but does not enable walking, limiting the type of exercise for people without access to pools. Lower-body positive pressure in itself has beneficial features, most studied with anti-thrombotic inflatable leggings and anti-shock trousers, but also in baro-physiology enhancing cardiorespiratory and autonomic nervous system function modulating molecules related to exercise and the chronic inflammatory insulin resistant syndrome of obesity and its comorbidities [Supporting Table S1B].

We designed a brief (30 minutes), low exertion treadmill exercise program with lower-body positive pressure and infrequent sessions (2-3/week) lasting 10-12 weeks in a sedentary obese female population not inclined to lose weight for cultural reasons [49], with limited access to pools and gyms living in neighborhoods with unsafe streets and parks.

## Methods

### Subjects

Volunteers among hospital employees and their families (not related to or supervised by the investigators) were recruited through word-of-mouth at SUNY Downstate Medical Center being offered participation in an exercise study of “metabolic fitness”, defined as “the ability of the body to use energy from dietary sugars and fats”. Eligible were otherwise healthy overweight–obese (BMI 28-50), weight-stable, untrained, non-diabetic, pre-menopausal Black women aged 18 to 56 years [Supporting Table S2], willing and able to commit to 2-3 times weekly 30-minute bouts of exercise on the weight-supporting treadmill for 12 weeks (more than 20 bouts) comfortably using the treadmill. They were not compensated but received a thorough metabolic work-up described below, signing consent forms covering all aspects of this study approved by the local institutional review board. All assays and measurements, treadmill supervision and statistics were performed by investigators blinded to accrued phenotypic data.

Exclusion criteria were shift-work and strenuous work-related functions, participation in dieting or exercise programs the last 3 months, surgical treatment of obesity, being pregnant or planning pregnancy, being smokers, taking medications known to affect energy balance or having musculo-skeletal or other conditions incompatible with using the treadmill. We excluded subjects with known diabetes, hypertension and dyslipidemia and those using psychotropic medications potentially affecting appetite regulation, oral contraceptives or steroids of any type. They consumed no or only minimal alcohol.

### Baseline history, physical and activity assessment

A structured 45-50-minute interview based on questionnaires exploring demographic and socio-economic factors, food insecurity, household stressors and a thorough medical history was conducted. Anthropometric measures included weight, height, BMI, waist and hip circumferences. Blood pressure and heart rate were measured after subjects had been seated for 5 minutes. Taking into consideration the cultural habits and beliefs of our population, there was no “weighing in” or mention of body weight, diet or dieting during visits to the laboratory. Most subjects were administrators engaged in typically sedentary work. Daily activity was exclusively measured pre- and post-study using a wrist-worn Fit-Bit^®^ accelerometer for two 3-day periods one of which included one non-working week-end day [Supporting Methods].

### Fasting morning blood and oral glucose tolerance tests

An in-dwelling ante-cubital venous catheter was placed with the patient seated in a comfortable recliner. At time 0 of a standard 75 g oral glucose tolerance test (OGTT) blood was drawn for determination of glucose and insulin, HOMA-IR, GLP-1, GIP, free fatty acids (FFA), C-peptide, fasting C-reactive peptide (CRP), Interleukin-6 (IL-6), TNFα, leptin, total ghrelin, total adiponectin, GIP, glucagon, triglycerides (TG), HDL-cholesterol, oxytocin (OXT), β-endorphin and orexin A (ORA). During OGTT at 15, 30, 60, 90 and 120 minutes blood was drawn for glucose, insulin, C-peptide, GLP-1 and FFA, allowing calculations at 2 hours and area under the curve (AUC). Blood chemistry details are provided in Supporting Methods.

### Weight-supporting treadmill

The lower-body positive pressure (LBPP) treadmill (AlterG^®^) [ref.50] consists of a stationary heavy metal frame with a movable gantry that can be raised to the level of a user’s waist and is attached to an inflatable clear-plastic malleable “skirt” surrounding the electric treadmill. [Supporting Figure S1] The waist of the skirt is encircled with one portion of a zipper corresponding to a zipper around the waist of neoprene shorts, worn by the subject. The treadmill’s mechanism covertly measures the person’s weight, enabling determination of a percentage of body weight to be off-loaded by treadmill pump inflation increasing pressure on the lower body. The maximum pressure exerted is equal to that of standing in water in a pool (~ 1 psi; 51.7 mmHg; 70 cm H_2_O; 6.9 kPa) [ref.51].

### Exercise protocol

Off-loading was arbitrarily set at 40% reduction of body weight, subjects thus exercising on the treadmill at 60% of their weight throughout the study. Speed set by the subject allowed completion of a 30-minute session. Initial incline was set at 0°. Subjects were encouraged to increase the speed with time and staff gradually increased the incline every 3 sessions to 3, 5, 7, and a maximum of 10%. Time, distance, speed, treadmill incline and % weight offloaded at the 25-minute mark was recorded after each session. After completion of 20-24 sessions OGTT with fasting blood tests similar to before the exercise regimen were performed more than 24 hours after the last session.

**Energy expenditure** We calculated metabolic equivalents (METs) of energy expenditure according to standard equations [52], but adjusted for weight in our study of weight-supported exercise. We used the Harris-Benedict Equation for RMR to improve the accuracy of MET estimates to determine “corrected METs” when offloading body weight with the LBPP treadmill [53].

Corrected METs, kilocalories (kcal) and Watts (W) expended per session according to ACSM formulas, total exercise per week as MET minutes and kcal expenditure were used to compare our exercise intensity at 60% body weight with CDC and ACSM exercise recommendations at full body weight [54] adding up total energy expended by each subject during all sessions.

### Statistics

Means and standard deviations were compared by standard Student t-tests or Wilcoxon’s rank-sum tests based on a one-sample Kolmogorov-Smirnov test against a normal distribution pdf. We used one-way ANOVA to compare pre- and post-exercise effects and calculated Pearson or Spearman correlation coefficients between relevant metrics. Pre-post differences were evaluated using 2-tailed tests throughout, although our biomarkers had ample peer-reviewed literature supporting directionality of our findings [Supporting Table S1A], which would justify one-tailed tests. Statistical analyses used SPSS version 23 (SPSS, Chicago, IL).

## Results

### Baseline characteristics and proteomics

According to our inclusion and exclusion criteria and the characterization in Supporting Methods and Table S2 these 16 subjects were overweight/obese and sedentary but were not diabetic, dyslipidemic or hypertensive. There were no statistically significant correlations between BMI and baseline molecules. In addition to traditional diabetes and dyslipidemia biomarkers we analyzed an array of incretins, inflammatory cytokines, adipokines and neurometabolic peptides individually known to be responsive to exercise and lower-body positive-pressure [Supporting Tables S1A and S2A]. We found novel correlations relevant to our post-study outcomes [Tables 1A and 1B].

**Table 1.**
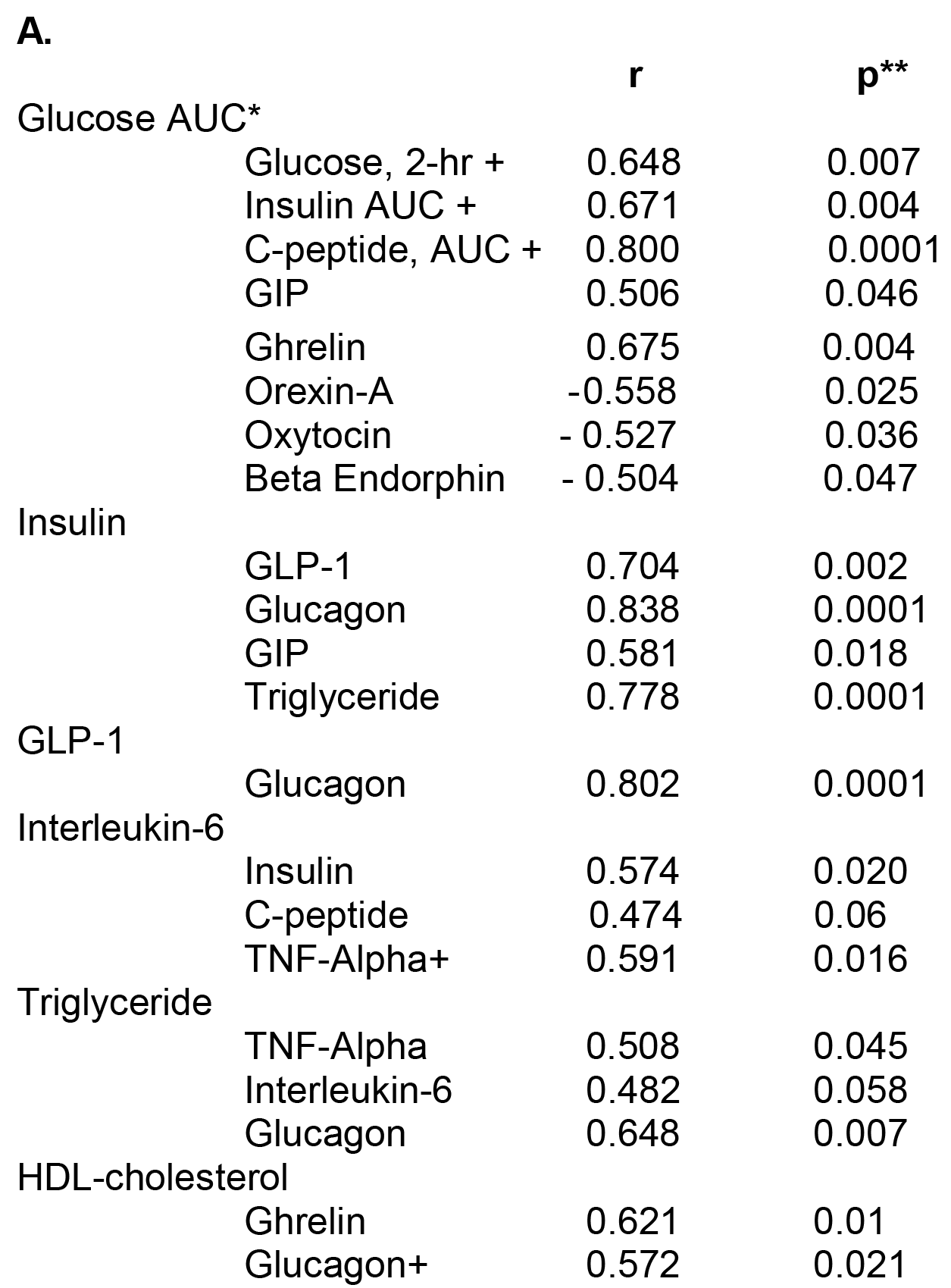
Baseline correlations of plasma cardio- (A.) and neuro-metabolic and appetitive (B.) molecules.

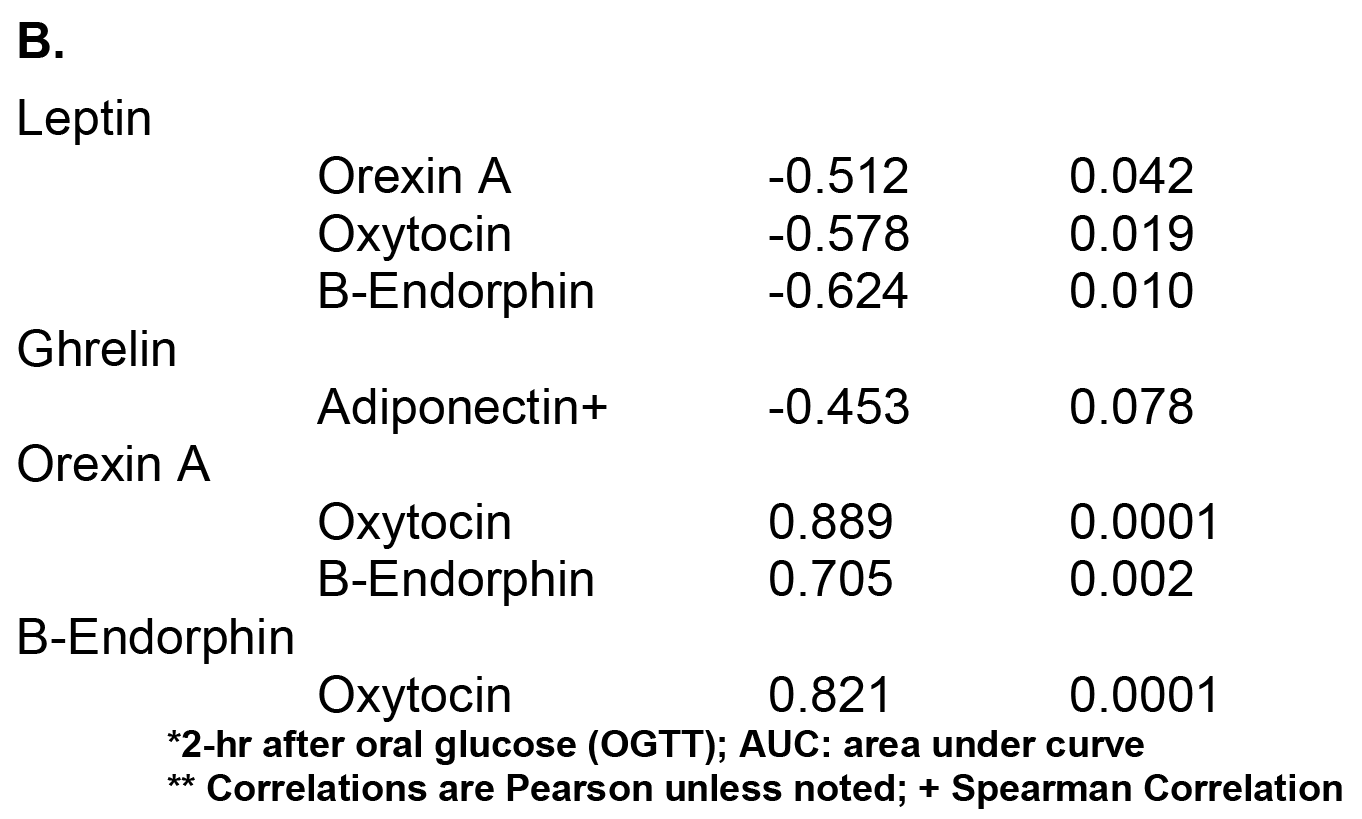

### Exercise and activity

Subjects exercised for 30 minutes 2-3 times weekly (mean 2.2) for 11 weeks on the Alter-G treadmill at 60% body weight without complications. They walked/jogged a mean distance of 2 miles (3 210 m) each session at a mean rate of 4.3 mph. The weekly addition of 4 miles corresponded to a 19% increase in weekly ambulation from 21 to 25 miles. The average level of exercise intensity was 8.5 METs calculated as corrected METs at 60% body weight, representing the lower end of recommended weekly expenditure [Table 2: ref. 53].

**Table 2.**
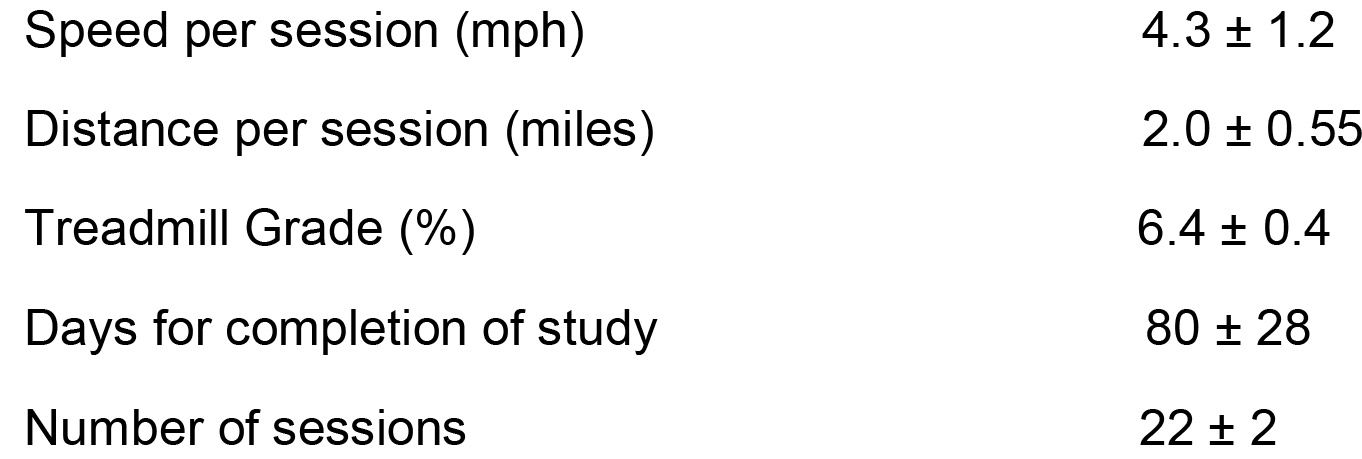
Exercise and energy expenditure during twice-weekly LBPP treadmill sessions (mean ± SD)

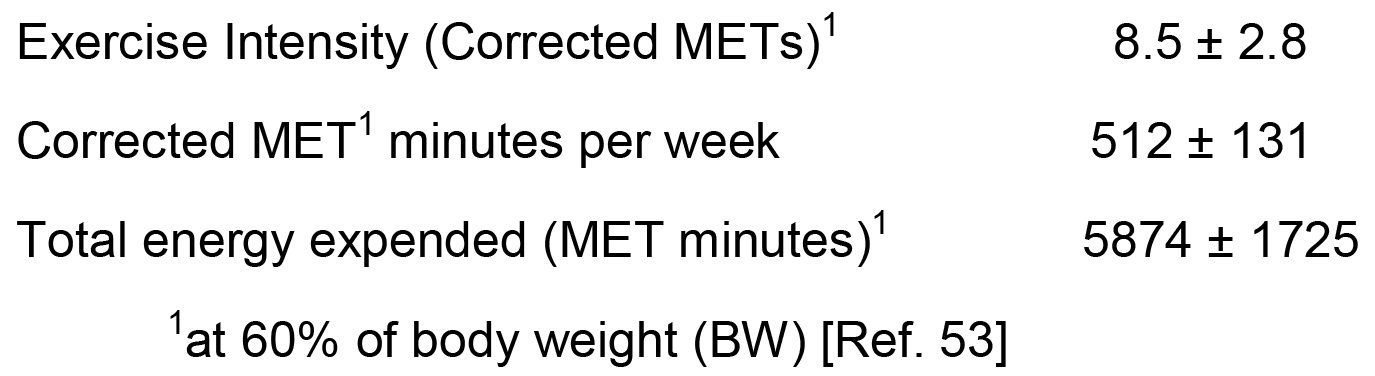

### Cardio- and neuro- metabolic changes

There were no statistically significant before-after mean differences in body weight, waist circumference or blood pressure. Eight subjects gained (range: 1.0 – 7.0 kg) and 6 lost weight (range: 2.2 – 9.5 kg), two exhibiting no change. Subjects, who had lost weight, disclosed *post hoc* during informal follow-up interviews that they had on their own dieted during and for several months following the study. Nevertheless, there were no statistically significant differences in outcomes between those with weight loss or weight gain. The mean duration between last exercise bout and blood testing was 9.8 days (range 1 - 44 days), with no statistically significant relationships between time post last bout and any outcome described in the following.

#### Dynamic Metabolic Testing (OGTT)

The 75-gram oral glucose challenge showed marginal decreases in fasting 2-hour and incremental *glucose* area after exercise, whereas both 2-hour [Figure1] and incremental *insulin* responses decreased significantly (34 and 15 % respectively), in agreement with a significant 15% decrease in 2-hour C-peptide and 19% decrease in 2-hour GLP-1 (p=0.06) [Table 3], together supporting reduced insulin resistance.

**Figure 1.**
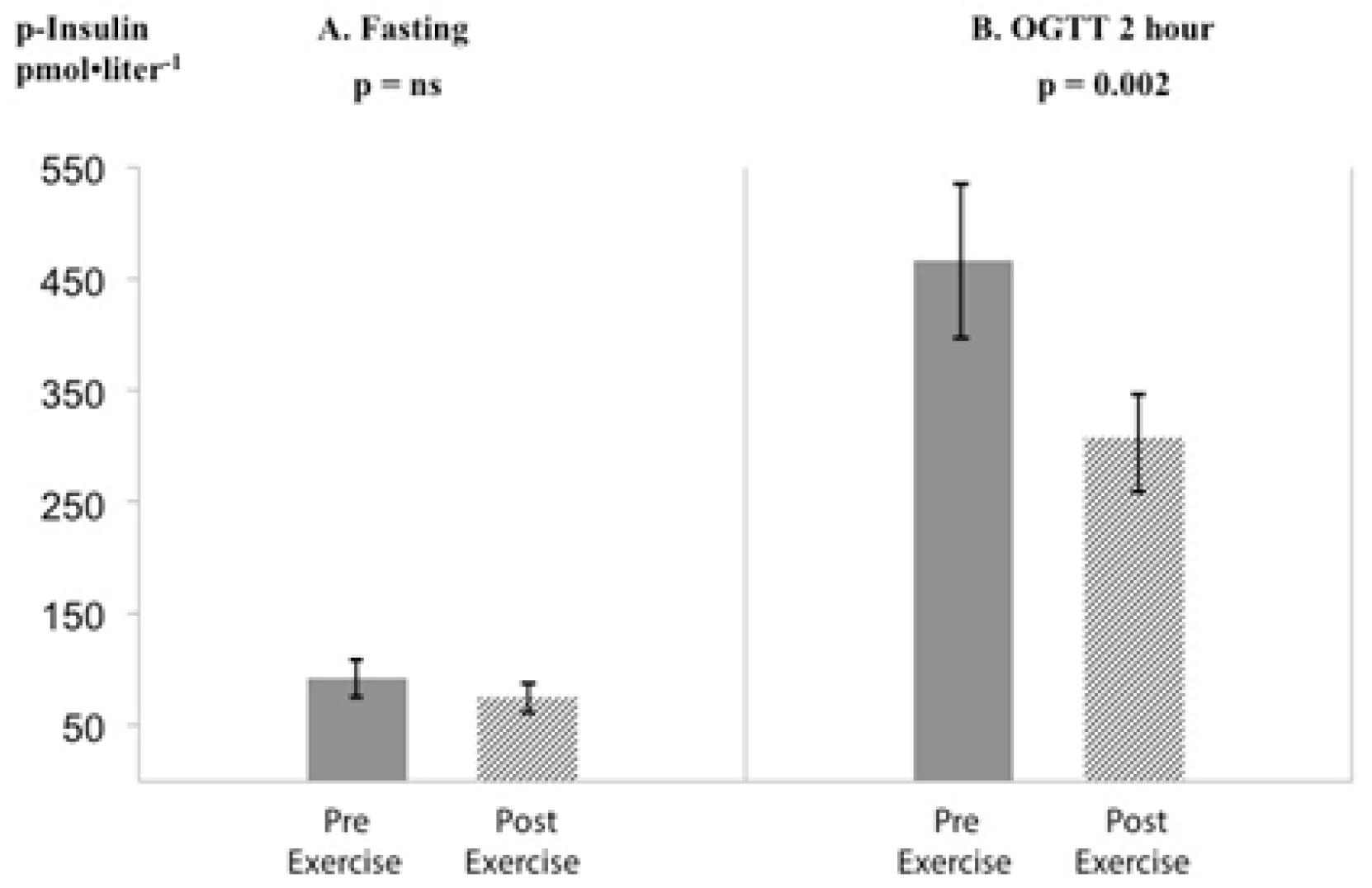
A. Fasting p-insulin and B. OGTT 2-hour p-insulin pre- and post-weight supported exercise (n=16; mean ± sem).

**Table 3.**
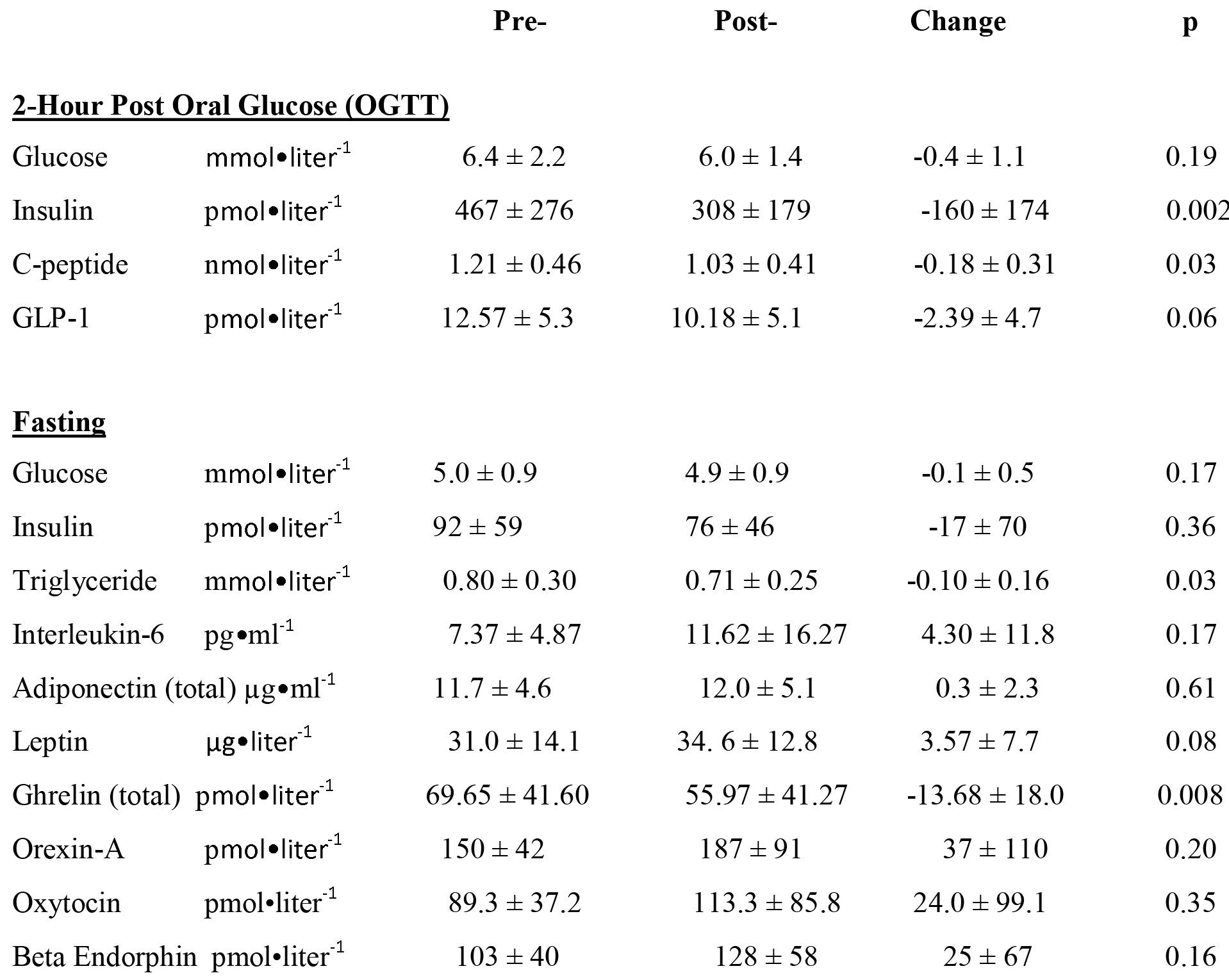
Changes in plasma cardio-, neuro-metabolic and appetitive molecules before (Pre-) and after (Post-) bodyweight-supported exercise (mean ± SD)

#### Fasting cardiometabolic molecules

Several differences in fasting plasma parameters were noted comparing before to after exercise. Glucose in the normal range decreased slightly (p=0.17) while there was a 19% reduction in fasting plasma insulin [Figure 1] with a marginal increase in adiponectin (2.6%) and decreases in C-peptide, GIP and glucagon. HDL cholesterol increased minimally whereas cortisol was virtually unchanged. Triglycerides, not elevated in any subject at baseline, decreased 11% [Figure S2A] with discordant changes in free fatty acids; leptin increased 12 %, (p= 0.084).

Changes in inflammatory cytokines varied between subjects according to base-line status, undetectable in some, divergent in others. C-reactive protein (CRP) elevated in 7 subjects decreased by 25% (p=0.052) and TNFα levels decreased in 7 subjects with detectable levels pre-exercise. Anti-inflammatory Interleukin-6, detectable at baseline in 8 subjects (mean 7.4 ± 4.9 pg/ml) increased 58% post study (mean 11.6 ± 16.3; p=0.170).

#### Fasting neuropeptides

Plasma ghrelin had decreased by 20% at the end of the study (p<0.008; Figure S2B) whereas Orexin A increased 25% (p=0.169), oxytocin 27% (p=0.348) and β-endorphin 24% (p=0.156). These latter relatively large within-subject increases, although statistically insignificant by 2-tailed t-tests owing to the relatively small numbers of subjects, were also assessed for ‘responder’ vs. ‘non-responder’ status, as well as by median split exhibiting statistically robust differences (all p<0.01, not shown), consistent internally and with existing exercise and baro-physiology literature.

#### Correlations between before-after changes

There were no statistically significant correlations between body weight change or total energy expended (MET minutes) and decreases in OGTT glucose, 2-hr insulin, C-peptide, fasting triglycerides or ghrelin. Significant correlations were present between total MET minutes and decrease in glucoregulatory 2-hr GLP-1 (r = − 0.679; p= 0.004) and the marginal increase in adiponectin (r= 0.471; p= 0.066). Changes in 2-hour GLP-1 were unrelated to changes in appetitive peptides.

Although parametric statistical analyses did not detect significant before-after changes in many peptides the changes were remarkably highly and consistently correlated. Intercorrelations between drops in gluco-regulatory 2-hr glucose, related to TNFα, and 2-hr insulin and C-peptide were significant; 2-hr insulin, in turn was highly correlated with rises in β-endorphin, orexin A and oxytocin [Table 4]. Baseline levels of leptin, marginally correlated with 2-hr insulin (r= 0.38; p=0.143), predicted the reduction in 2-hr insulin (r= 0.566; p=0.022). Significant correlation between increases in fasting leptin and adiponectin (r= 0.718; p=0.002) [Figure 2] reflected improved insulin sensitivity concordant with significant reductions in insulin and C-peptide. HDL-cholesterol change correlated with FFA AUC, leptin and adiponectin but not with any inflammatory cytokines [Table 4].

**Figure 2.**
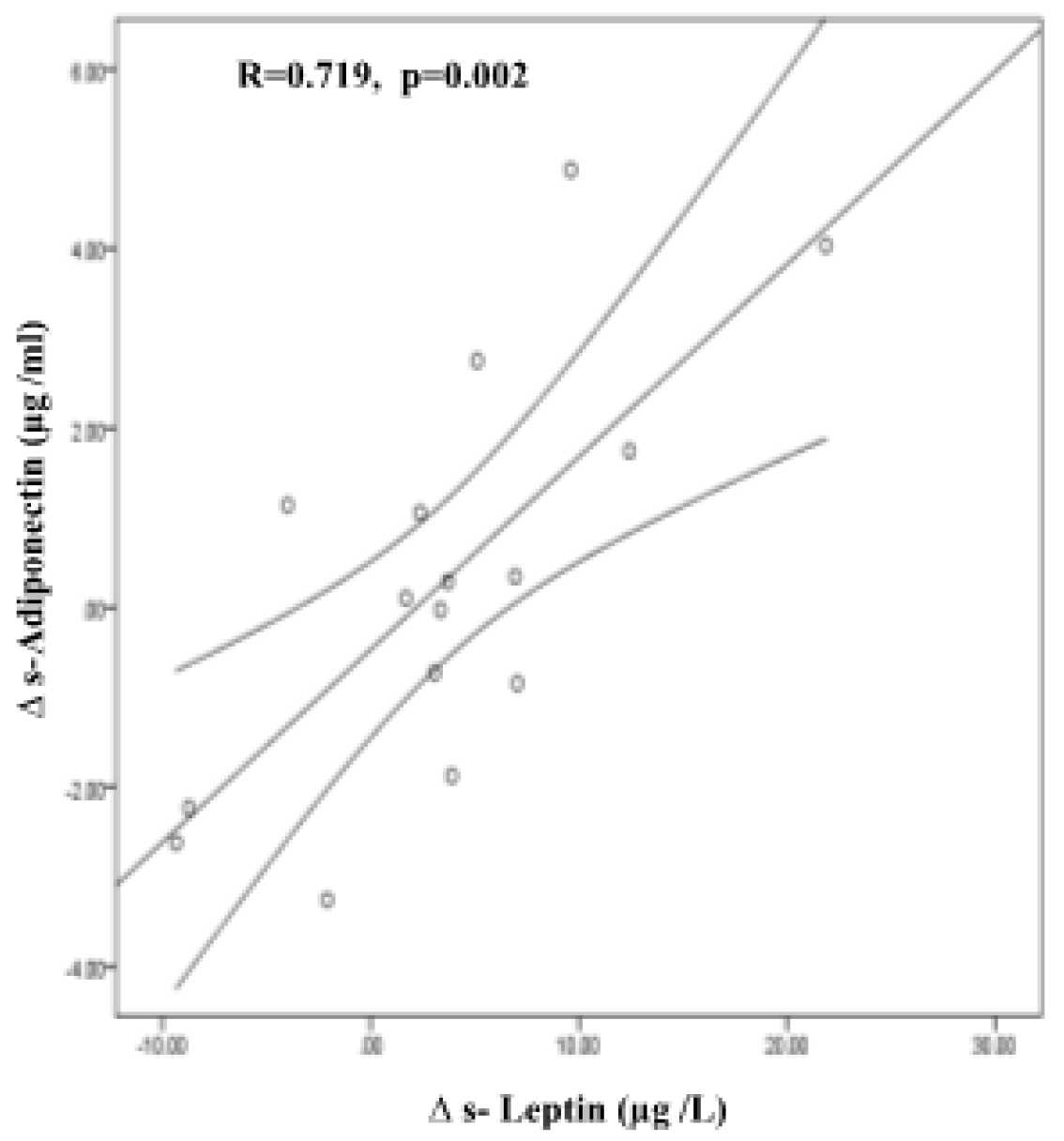
Pearson correlation of changes (Δ) in s-adiponectin and s-leptin, pre- and post-weight supported exercise. [Y=0.47+0.21X; r=0.719; p=0.002. Fit line 95% CI for the mean. (n=16)].

**Table 4.** Correlations between changes in cardio- and neuro-metabolic molecules.

|  | **r** | **p\*** |
| --- | --- | --- |
| **Glucose, 2-hr\*** |  |  |
| Insulin, 2-hr* | 0.445 | 0.084 |
| TNF $\alpha$ <sup>+</sup> | 0.631 | 0.009 |
| Leptin | 0.482 | 0.059 |
| Ghrelin | 0.457 | 0.076 |
| Orexin A | -0.502 | 0.048 |
| Oxytocin | -0.422 | 0.103 |
| Beta Endorphin<sup>+</sup> | -0.517 | 0.037 |
| **Insulin, 2-hr\*** |  |  |
| C-Peptide, 2-hr<sup>+</sup> | 0.538 | 0.031 |
| Triglyceride | 0.357 | 0.174 |
| Leptin | 0.475 | 0.063 |
| Orexin A | -0.594 | 0.015 |
| Oxytocin | -0.690 | 0.003 |
| Beta Endorphin | -0.487 | 0.056 |
| **C-Peptide, 2-hr\*** |  |  |
| C-Reactive Protein | 0.451 | 0.079 |
| **GLP-1, 2-hr\*** |  |  |
| Glucagon | 0.555 | 0.026 |
| IL-6 | 0.462 | 0.071 |

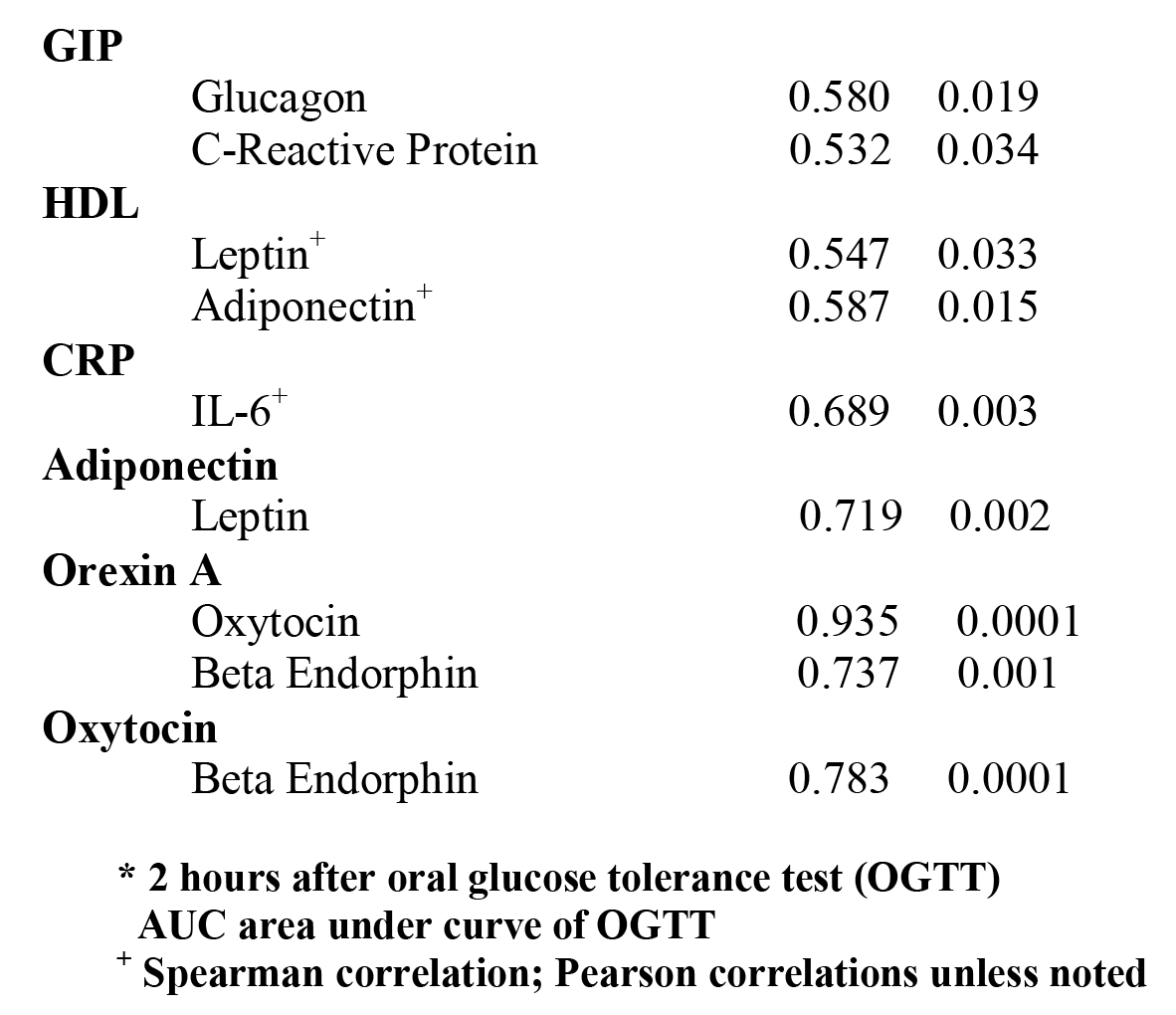

We found previously un-reported highly significant correlations between before-after exercise changes in *neuro-endocrine* peptides in agreement with the pre-exercise correlations and unrelated to total energy expended, weight changes or any phenotypic data [Table 4].

Altogether these findings demonstrate consonant before-after reductions in gluco-regulatory molecules reflected in correlations between glucose, insulin, C-peptide and incretins during OGTT. These reductions correlate with inflammatory markers, adipokines and neuroregulatory, appetitive peptides.

## Discussion

### Exercise

#### Cardio-metabolic factors

The preliminary findings of this unique study challenge the premise of the preponderance of studies and guidelines for treating and preventing diabesity, viz. that body weight is of primary importance for the disease and requires intermediate to high levels of frequent physical activity to improve performance and to cause weight loss [55]. Our sedentary subjects did not lose weight from the low level of exertion in the sedentary range and, indeed, there were no associations between energy expended or weight loss or gain and improved cardio-metabolic metrics such as insulinemia, glucose tolerance markers and plasma triglycerides earlier described in literature supporting diet and exercise guidelines.

Robust reductions in plasma triglycerides in the normal range in our Black subjects, normally with lower levels than those with other geographic ancestry [56, 57], are significant in the context of increased recognition of low-moderate triglyceride levels as cardio-metabolic risk factors [58] furthermore supporting the need to redefine blood lipid standards for Black people [57]. Hyperinsulinemia has long been known as a driver of hepatic triglyceride synthesis [59], which accords with our present finding of concomitant reductions of OGTT insulin and fasting plasma triglycerides after our low intensity exercise which otherwise is found after more intensive exercise [60, 61]. Our findings regarding *inflammatory markers* are similar to those of other studies of obese populations, although we generally excluded subjects with advanced disease sufficient to manifest detectable elevations of cytokines measured by routine methods. Thus 7 subjects had detectable elevated CRP and 7 elevated TNFα, all of whom reduced their levels post study. Three subjects did have impaired glucose tolerance (IGT) which normalized post-study independent of weight changes. These three were among the subjects that increased post-study levels of the anti-inflammatory IL-6, known to be beneficially elevated by exercise [9, 62].

#### Neuro-endocrine peptides

Remarkably the low amount of low intensity exercise combined with mild lower-body positive pressure was associated with effects on a wide array of cholinergic regulatory central and peripheral peptides. There was a substantial reduction in fasting levels of orexigenic ghrelin, a rapid sensor of fluctuations in nutrient stores which was not associated with the borderline increases in ‘satiety’ peptides leptin and β-endorphin or orexin A shown in others’ exercise studies [Supplement Table S1A]. Since glucose is the primary substrate in brain, skeletal muscle and whole-body metabolism our finding of a strong relationship between the reduction of glucose area under the curve and ghrelin reduction (p<0.004) is consistent with the primary response to exercise preceding transition to lipid and amino acid utilization in the sedentary range. Also there were no relations between ghrelin and glucose-stimulated levels of the pleiotropic neuro-endocrine brain-gut peptide GLP-1. Although it is synthesized in the ileum and mediates incretin effects associated with insulinotropic pancreatic β-cell secretion exhibited in this study, GLP-1 is also known for its central effects suppressing appetite and increasing neurogenesis [9, 63], here demonstrating discordance between its peripheral and central effects, possibly related to improved GLP-1 receptor function.

The robust correlations between leptin and β-endorphin, orexin A and oxytocin and among these latter peptides are intriguing, given their respective roles in energy balance, neurogenesis and mood [64, 65, 66]. Although not the subject of this study, documented beneficial effects of exercise on anxiety and mood disorders [67, 68] might be expected, related to our neuro-peptide changes after low-intensity exercise, especially in the absence of pain, a powerful mediator of mood. Informally all of our subjects have later spontaneously expressed their enjoyment of this exercise, especially those with experience with conventional treadmills. The brain-gut peptides in this study participate in cholinergic circuits which modulate appetite suppression on downstream targets in the hypothalamus [69] and also affect pancreatic structure, function and release of neurotrophic peptides other than GLP-1.

Our heterodox findings of improved glucoregulation with triglyceride decrease, mild effects on cytokines and adipokines but a robust effect on the orexigenic peptide ghrelin in the absence of weight loss accompanying brief, low-intensity exertion, raises questions about putative mechanisms. We speculate that improvements in substrate utilization such as glucose disposal, lipolysis or “protein sparing” are unlikely to be primary determinants but rather synergistic with autonomic nervous system activation/conditioning (“stress buffering”) [2] and/or release of key molecules through mild compression and use of lower extremity muscles known to have beneficial cardiovascular [42, 43], angiogenic and metabolic effects associated with restored autonomic balance.

### Baro-physiology

Standing, *per se* improves cardiometabolic risk when used as a break in sedentary activity [70] as do exercise “snacks” [71]. The compression exerted by the lower-body positive pressure treadmill during 40% off-loading is similar to standing in a swimming-pool unrelated to muscle use. We have demonstrated decreased heart rate during weight offloading during standing [44], recently shown by others during increasing levels of exertion during LBPP [72]. We speculate that there is synergy between standing and positive lower body pressure which would cause local hypoxia increasing muscle perfusion and oxygen uptake [73, 37] as in pre-conditioning, thus potentiating the effects of the low intensity exercise improving mitochondrial function. Together with the described effects on the different classes of peptides we posit that lower-body positive pressure increases parasympathetic tone in the cranio-sacral division of the autonomic nervous system, also reducing dysautonomia. Although not studied in these subjects ample evidence supports effects of LBPP on sympathetic tone, hemodynamics, oxygenation and metabotropic molecules [Table S1B].

### Public Health Perspective

A recent review of obesity identifies “decreasing time spent in occupational physical activities and displacement of leisure-time physical activities with sedentary activities” contributing to epidemic obesity and “lack of effective and accessible life-style programs that can be administered locally or remotely at low cost to diverse populations” explaining why “only a fraction of patients for whom…treatments are indicated actually receive them” [74].

Our findings imply that it is possible to improve metabolic fitness in underserved populations with high prevalence of diabesity, barriers to exercising and cultural resistance to losing weight. The study coincides with reports of adverse effects of outdoor exercise in inner-city environments [75]. The indoor exercise in this study is provided by a safe and convenient weight-supporting lower-body pressure treadmill proven effective for ambulation in orthopedic and neurologic rehabilitation.

Effects of this magnitude on hyperinsulinemia, plasma triglycerides and ghrelin are exhibited after more time-consuming and intense volitional life-style changes or after metabolic operations recommended for type 2 diabetes [76]. The cost of an anti-gravity treadmill is well within the range of these modalities, requires substantially less personal investment of time to use and must be considered cost-effective since a single treadmill can be used by many subjects (e.g. in a family, therapeutically or preventively) over a long period of time, thus providing return on the initial investment regardless of payer.

### Limitations

Limitations are having a *selected population* of volunteering, obese, *female* hospital workers of Caribbean-Black ancestry not allowing generalization to other populations with different BMI standards and metabolic metrics. However, regardless of geographic heritage our subjects are reproductive-age obese, relatively low socio-economic status (SES) women, representative of a large stratum of prevalently obese populations.

Our *variances* are substantial, reflecting “real-world” clinical research in our environment with significant confounders, yet achieve statistical significance with trends consistent with exercise studies in very different populations and designs (from trained Caucasian male athletes to exhaustive sub-maximal acute studies). We were unable to coordinate pre- and post-exercise blood sampling with our subjects’ *follicular phase*, which likely also has contributed to the variance. Time of day was variable for practical reasons. We did not detect any seasonality; the exercise was performed in a temperature-controlled gait laboratory, but we did not measure body temperature.

By design we omitted metrics of diet (macronutrient distribution, 3-day diaries), appetite (hunger ratings, taste preference), size and body composition (weight, BMI, lean body mass) and physical performance (O_2_ consumption, lactate production) leaving these to future studies. Thus, our study is exploratory and mechanisms remain to be fully determined, although many of the findings are hypothesis-generating.

## Conclusions

We conclude that weight-supported lower-body positive pressure enables low intensity ambulation in subjects otherwise not inclined to exercise who hereby experience beneficial cardio- and neurometabolic effects without pain. This level of energy expenditure, in the sedentary range, is unlikely to trigger counter-regulatory mechanisms induced by glycogen depletion common to moderate-high intensity exertion of longer duration recommended in public health guidelines for weight loss and prevention of diabetes and co-morbidities. Amelioration of the dysautonomia of allostatic load is a significant explanatory candidate mechanism, supported by numerous pre-clinical and clinical studies of effects of exercise and lower-body pressure stimulating the parasympathetic nervous system, modulating cardiac autonomic function [77], central neurogenesis and cognitive function [78, 79] associated with mood.

## Acknowledgments

We are grateful to our many students and research fellows who participated in recruiting and supervising participants, setting up accelerometer studies and conducting interviews. They appear as authors in our published meeting abstracts. Stephen McCormick, MD, MPH graciously assisted with bio-statistics during early stages of data analyses, which we appreciate. Thanks also to our volunteer participants who found time before and after work and enthusiastically recruited by word-of-mouth. This study was made possible through a generous unrestricted equipment loan from AlterG, Fremont, CA, who had no part in the funding, design, interpretation or writing of this study.

---

**Figure S1.**
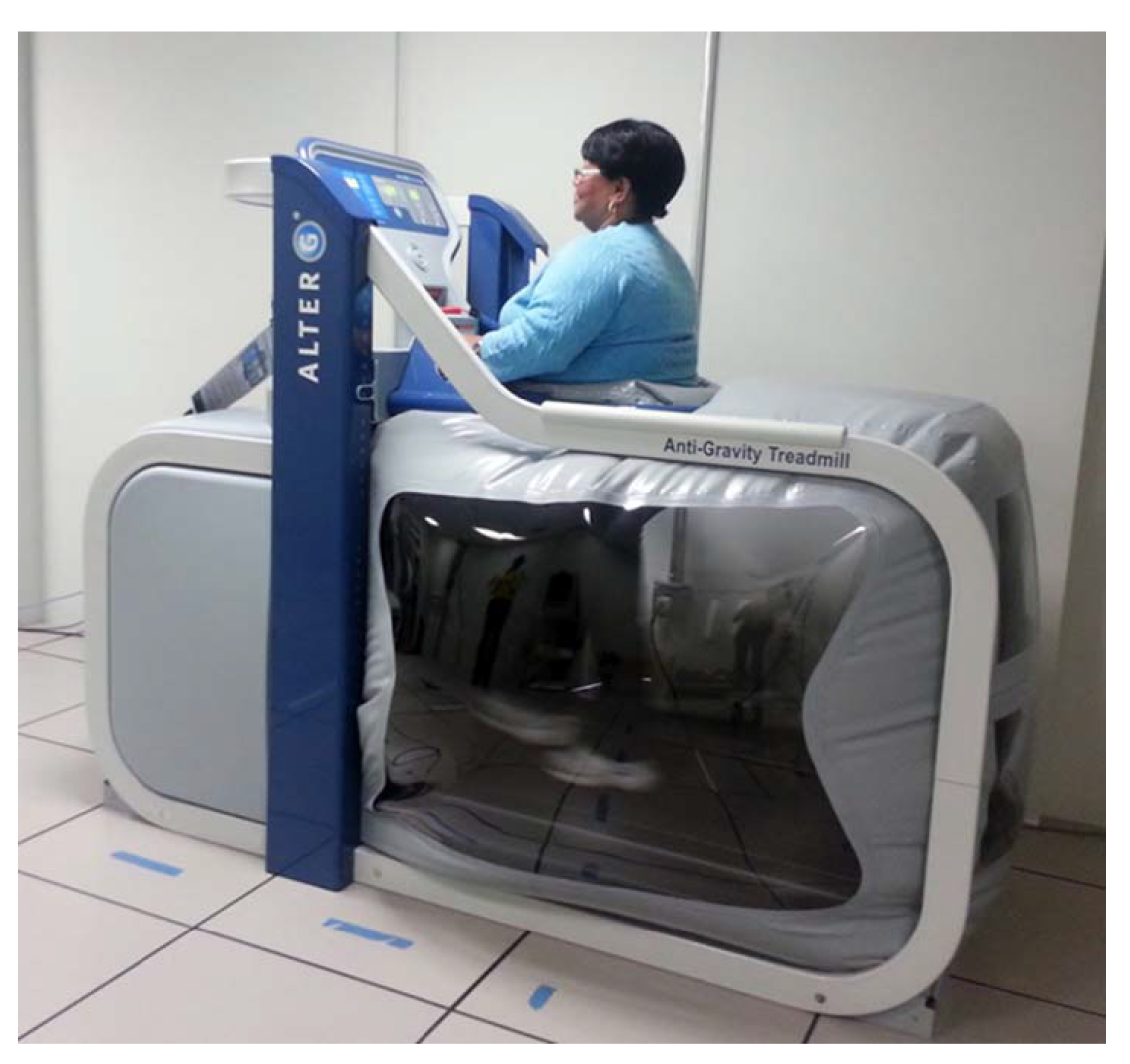
Lower-body positive pressure treadmill (AlterG)

